# Elongated sample plot shapes produce higher measurement precision in clustering populations

**DOI:** 10.1101/2021.09.10.459832

**Authors:** Richard McGarvey, Paul Burch, Janet M. Matthews

## Abstract

Monitoring the density of natural populations is crucial for ecosystem management decision making and natural resource management. The most widely used method to measure the population density of animal and plant species in natural habitats is to count organisms in sample plots. Yet evaluation of survey performance by different sample plot shapes, e.g. quadrats compared with transects, has been largely neglected since the 1990s and has not been undertaken using simulation. Simulating populations and surveys, we evaluated population density measurement precision for 900 cases, testing 30 sample plot survey designs in each of 30 spatially clustered populations. We varied three design options: elongation of plot shape while keeping sample area constant, systematic or random plot allocation, and sample size. Survey design performance varied markedly: elongating the plot shape always improved survey precision; allocating plots systematically sometimes did. (i) Averaged across all tested populations, elongated (1:100) transect plot shapes were 2-to-3 times more precise than square (10:10) quadrats. (ii) The precision of systematic surveys accelerated with sample plot number, increasing faster than the (known) linear increase under simple random sampling. This non-linear, concave upward, dependence of systematic precision on sample size has not previously been reported. (iii) The most precise design we evaluated used long narrow transects allocated systematically. Averaging among all 30 tested populations, a researcher would need 600 random square (10:10) quadrats to equal the precision achieved by 100 systematic (1:100) transects. Finding this average efficiency difference of 600% for a survey sample size of 100 plots, these simulation results imply that field trips requiring five sampling days using random quadrats could achieve equal precision in one or two days using systematic elongated transects. For all clustered populations we tested, long narrow transects resulted in a more efficient design for sample plot survey.

## Introduction

Under climate change, species introductions, habitat displacement, and other human impacts, native populations worldwide are threatened. Monitoring this change is crucial but expensive, and resources are limited. Survey efficiency is critical. Yet, less effort has been devoted in recent years towards improving the conventional method of measuring the density of natural populations. Statistical methodology of survey design has shifted towards adaptive sampling and distance-based techniques, mathematically sophisticated approaches that require complex field sampling protocols. But evaluating conventional survey methods, of counting organisms within allocated sample plots, offers a more direct avenue of progress for conservation monitoring, by improving the information content of standard population measurement data and reducing the number of fieldwork days required for its collection [1]. Sample plot survey designs offer intuitive simplicity in field implementation where navigating complex habitat and undertaking organism counts can offer challenge enough, and standard statistics can be applied to the resulting data. In 12 issues of Ecology reviewed (Articles, May 2016-April 2017), the majority of field studies measuring organism abundance employed sample plots, with 24 using random quadrats, 20 using systematic quadrats, 7 using random transects, and 15 using systematic transects. Among these, 44% took consideration of a habitat gradient.

A principal challenge to efficient measurement of population density by any method is the clustering of organisms in space [2, 3]. This nearly ubiquitous autocorrelation of organism locations in two-dimensional habitats reduces the precision of field sampling, for example, in fisheries and marine surveys [4-9], insects [10], plants [11], and in biodiversity assessment [12]. Population clustering increases the skewness of the sample distribution, typically yielding many zero counts [13] and a few very high values, widening confidence intervals of measured population density. In population change hypothesis testing, environmental impact analysis, or modeling, this often high error in measured population abundance obscures statistical inference. Our aim here is to evaluate the performance of a wide range of conventional sampling designs, and to identify those yielding highest precision for measurement of absolute population density, in a wide range of clustered populations.

To evaluate survey design precision, spatial simulation is now a standard tool. It is flexible, accurate and robust. First, one or more simulated spatial point populations are generated within a bounded study region, each point designating the location of one organism. Researchers control the forms and degrees of clustering in simulated populations through the spatial autocorrelation parameters of the population-generating algorithm. Second, simulated surveys measuring mean population density are run according to the specifications of each tested survey design. For this study, we allocated survey sample plots of given rectangular shape within a bounded study region containing clustered spatial point populations and simulated organism counts were taken. The precision of each design in each population is estimated from the variance of Monte Carlo-iterated survey density means. To date, perhaps surprisingly, simulation has not been used to compare the precision of different plot shapes. Here we address and extend this objective, using simulations to evaluate combinations of the three basic sample plot survey design options: plot shape, plot allocation, and sample size.

In early work, the precision of different plot shapes was investigated using completely enumerated field populations. Christidis [14], reviewing early studies for crop yields, recommended long narrow sample plot shapes to be more precise overall (while citing Lyon [15] and Mercer and Hall [16] who did not), and emphasized an orientation parallel to fertility gradients. Clapham [17] with flowering plants and Bormann [18] within a North American forest tested differently shaped rectangular sample plots and found strips to provide lower variance than square quadrats. Kenkel et al. [19] in review advocated elongated sample plots citing earlier agricultural studies [20, 21]. In subsequent field studies using diverse approaches, all measuring crop yield, the outcomes were less clear [22, 23], with two studies finding no benefit to an elongated sample plot [24, 25] and one reporting variation among species [26]. To date, consensus has not been reached about the precision implications of plot shape. Moreover, there has been little examination in natural populations which usually show far more clustering than crops. Since the 1990s, investigation of plot shape has waned leaving the question unresolved.

The absolute density of a population that varies over space is undefined unless the area covered by the estimate is exactly specified. Surveys to measure absolute density therefore require bounded study regions. Within this study region, sample plots are representatively (here randomly or systematically) allocated.

Because conventional designs using random or systematic quadrats or transects can assume uniform sampling probabilities, they are a priori unbiased for the sample mean [27]. This is, of course, an important advantage of conventional sample plot surveys. All the conventional designs we evaluate here being unbiased, only their precisions are needed to rank survey design performance.

Evaluating precision for 900 combinations of survey design and study population, roughly two orders of magnitude more cases than previous simulation studies, presented several clear trends informing the choice of an efficient sample plot survey design.

## Methods

We evaluated and compared precision for combinations of three sample plot survey design options: (i) plot shape, as rectangular plots of varying width:length but constant (100 m^2^) habitat search area, (ii) systematic and random plot allocation, and (iii) sample size as number of equal-area sample plots.

The spatial simulations to evaluate survey design precision were undertaken in three steps: (1) generating 30 spatial point populations, (ii) surveying each population using each of 30 survey designs, and (iii) analyzing the results, specifically the absolute population density sample means from 10,000 iterated simulation surveys, to compute the sampling error variance for each combination of design and clustered population. Precisions for some plot shapes, and sampling error variance ratios for pairs of selected cases were also computed to evaluate survey design performance.

### Generating spatial point populations

We simulated surveys in a square study region, nominally 1-km^2^, though isomorphic expansion or contraction of spatial scale would leave the design comparison results unchanged. Within this study region we generated 30 spatial point populations [28] that varied from spatially random to highly clustered. We generated clustering using a commonly assumed model, of isotropic exponential decline in correlation with separation distance, [2, 29, 30]. Exponential clustering is characterized by (1) the distance scale of spatial autocorrelation and (2) the maximum cluster density. These clustering properties were controlled by two exponential variogram parameters of function *gstat* [31] in the gstat R package [32, 33]. The *range* parameter specifies the separation distance beyond which population densities are effectively uncorrelated; *psill* (partial sill) determines the average squared difference between densities at well-separated distances (greater than the *range*). The *nugget* (effectively the measurement error) was set to zero for all 30 populations. Point location maps of all 30 test populations, expressing varying degrees of clustering, are shown in Fig 1. Enlarged 1-m^2^ resolution maps for 18 of these populations are presented in Appendix S1. The R code used to generate these populations is given in Dataset S1. The 30 test populations all have a mean density near 0.1, varying from 0.095 to 0.110 organisms m^-2^. This total population size of about 100,000 was achieved, for each choice of *range* and *psill*, by randomly generating multiple populations and selecting the one with a mean density closest to 0.1 m^-2^. Thus, we held mean density approximately constant among 30 test populations and varied the two parameters controlling the distance scale and intensity of spatial clustering.

**Fig 1.**
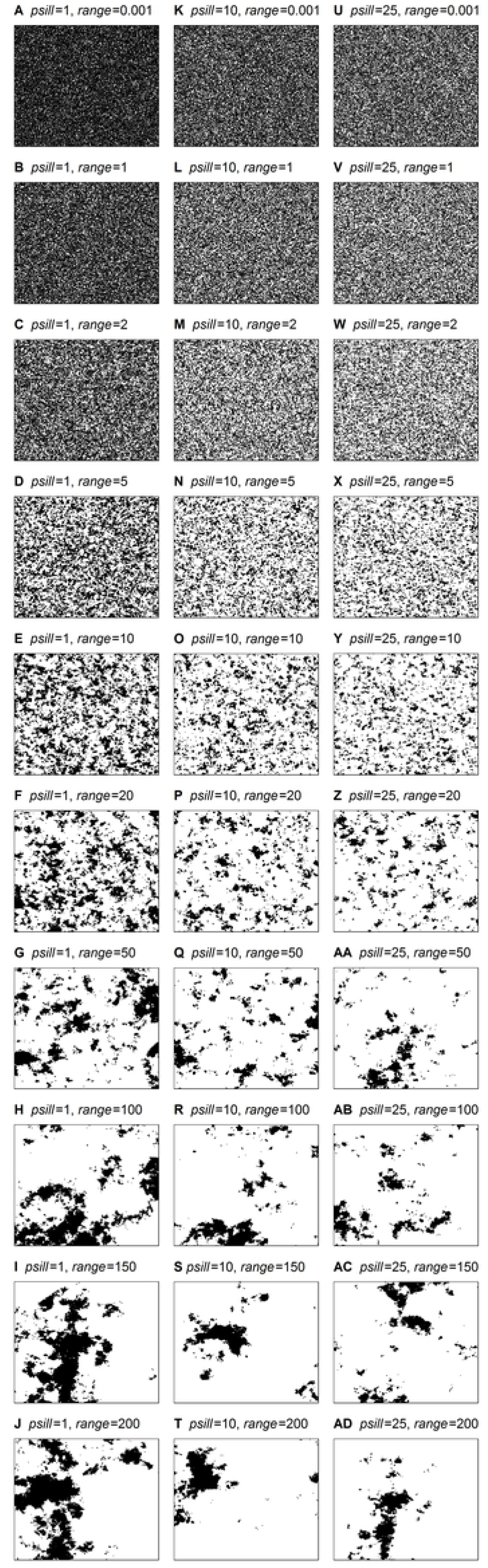
Thirty spatial point populations. Mapped point locations of organisms in the survey study region for each of the 30 spatial point populations used in simulation survey design performance comparisons.

### Simulating survey sampling

We simulated sample plot surveys by organism counts taken within rectangular plots allocated without replacement within the bounded study region of each spatial point population. We required that all sample plot shapes evaluated were of equal area, guaranteeing that all surveys (of given sample size) cover the same search area of habitat, and so have approximately the same survey cost. Lastly, we required that these equal-area rectangular sample plots provide a complete uniform coverage (tessellation) of the study region. This eliminates sampling edge effects or possibility of overlap in sample plots, removing these sources of uncontrolled variation from survey design comparisons. For each plot shape, 10,000 non-overlapping 100-m^2^ sample plots tessellate the 1-km^2^ survey study region, forming the sample frame. Geometrically, only certain combinations of plot shape and sample size permit tessellation. Within these geometric constraints, we extended a simpler survey simulation method [34] to test all possible combinations of sample plot shape, allocation, and number.

Five rectangular plot shapes, each covering a search area of 100 m^2^, satisfied these constraints. These five sample plots vary by their ratio of width:length from the longest most narrow transect of 1:100 (denoted by its length as ‘L100’), to 2:50 (L50), 4:25 (L25), 5:20 (L20), and lastly including a square quadrat of dimensions 10:10 (L10). The simulation density measure from each sample plot was computed without error from the number of enclosed organism location points.

Under a systematic design, one sample plot density measure is taken from each of *n* identical strata. Sample sizes (*n*) is this study were thus further restricted to those that (i) permitted the 1-km^2^ study region to be partitioned into *n* equal-area square strata, and such that (ii) each of the five rectangular sample plot shapes can tessellate each individual stratum. Three sample sizes of *n* = 4, 25, and 100 satisfied these constraints. For a systematic sample size of *n* = 100, the study area is covered by a 10×10 partition of 100×100 m strata, for *n* = 25 by a 5×5 partition of 200×200 m strata, and for *n* = 4 by a 2×2 partition of 500×500 m strata. A systematic sample of *n* = 16 plots, requiring strata of 250×250 m, could not be tessellated by three of the five tested plot shapes (L100, L25 or L20) and so was not used for most survey design simulation testing. For one objective, to evaluate precision versus sample size, we used only the plot shapes L50 and L10 that can also tessellate a 250×250 square stratum, permitting the inclusion of this fourth sample size, *n* = 16.

We tested the two standard methods of plot allocation, simple random and one-start aligned systematic [35, 36]. For simple random allocation, a survey sample of *n* plots was selected randomly, without replacement, from the sample frame. For one-start aligned systematic sampling, the standard, and simplest, version of systematic sampling, a fixed position for the single sample plot within each stratum was chosen at random and used in all *n* strata. The R code for simulated plot sampling is given in Dataset S2.

### Statistics and confidence intervals

Sampling error variance was computed as the unbiased sample variance of 10,000 means of population density from 10,000 numerically iterated surveys. Precision was computed as the inverse of the sampling error variance. Methods and results for estimating confidence intervals of the sampling error variances, precisions, and sampling error variance ratios are given in Appendix S2.

## Results

In Fig 2, we illustrate the simulation sampling scheme. For each of four survey designs we mapped one sample of *n* = 100 sample plots (in red). The clustering of the spatial point population surveyed (in green) is shown. Below each map of sample and population, the resulting histogram of 100 organism counts shows each design’s sampling distribution. More highly skewed distributions, with more zero and very high counts, are observed for (i) random compared to systematic plot allocation (left graphs versus right, Fig 2) and, to a greater extent, for (ii) a square quadrat plot shape compared with an elongated transect (top graphs versus bottom). More zero and high counts widen the sampling error variance (the standard error of the mean squared), reducing survey precision. Here we follow convention and define precision as the inverse of sampling error variance.

**Fig 2.**
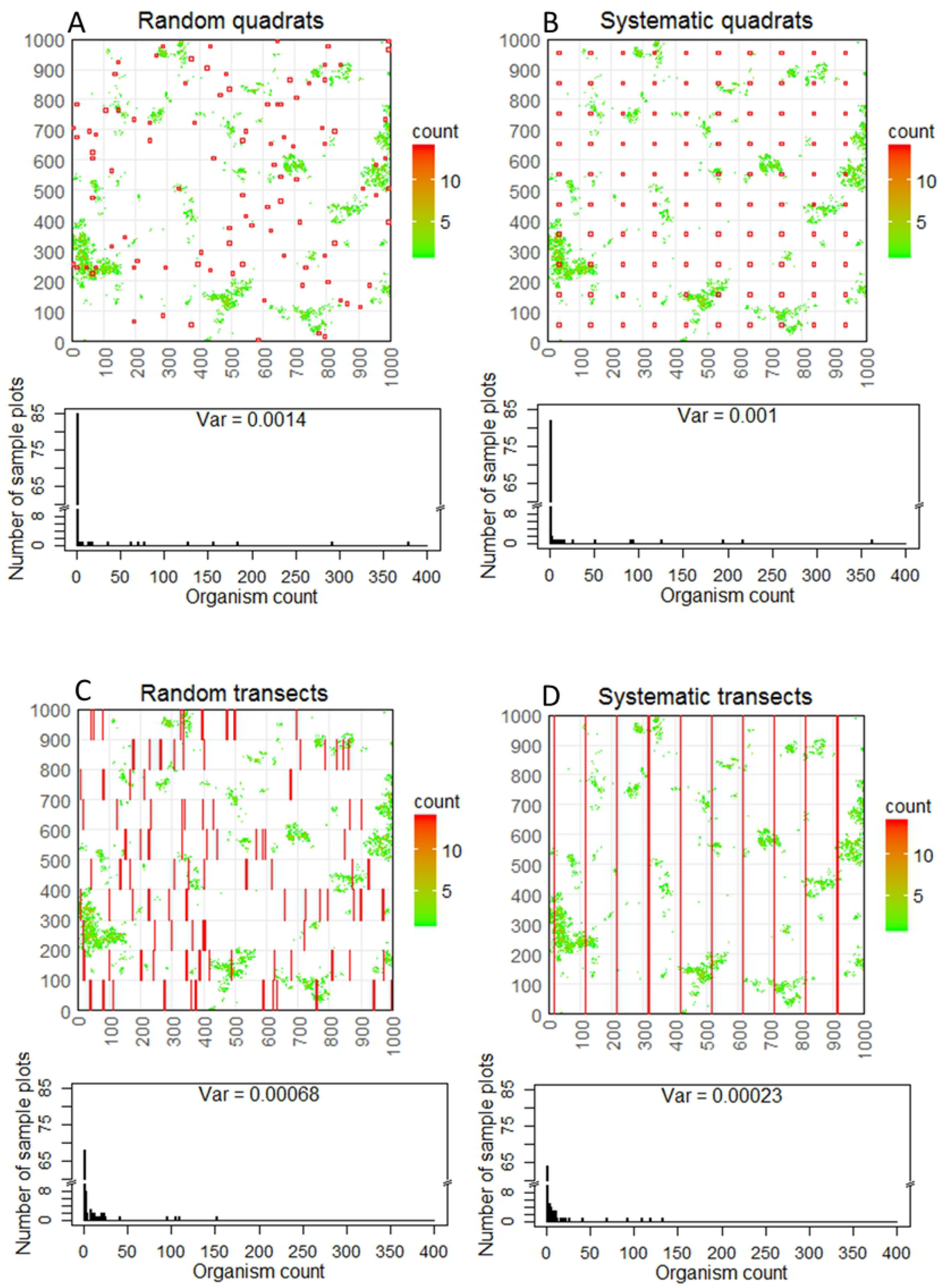
Samples drawn using four sample plot survey designs (*n* = 100) to measure population density. In our simulated 1000×1000 study region, four survey designs are illustrated: (A) random square (10:10) quadrats, (B) systematic square (10:10) quadrats, (C) random elongated (1:100) transects, and (D) systematic elongated (1:100) transects. Sample plot rectangles are drawn in red. The spatial point population sampled in all four panels (also mapped in Fig 1Q and Fig S1.10) was generated with variogram parameters of *range* = 50 and *psill* =10. Organism densities are color coded, higher densities in green, unpopulated areas in white. Because of the particular geometry of this 1-km square study region, 1:100 systematic transects oriented vertically span the full length of each (100×100) systematic stratum (Fig 2D), resulting in unbroken vertical sampling lines that span the survey region. These maps illustrate four of the 900 cases (four designs in one population) examined in this study. Below the map of each survey sample is its corresponding histogram of 100 plot counts, including the number of zero counts. The resulting sampling error variance for each case (“Var”) was computed as the variance of the 10,000 Monte Carlo-iterated survey estimates of population density. Each of these 100-plot samples shown was the first of 10,000 iterations drawn for each case.

For each of 30 survey designs (five plot shapes, systematic or random, three sample sizes) in each of 30 spatially clustered populations (three values of *psill* and 10 of spatial autocorrelation *range*), we computed the sampling error variance for the estimate of population density from 10,000 simulated survey density estimates. All 900 sampling error variances are plotted as bar heights in Appendix S3. Large differences in variance among the 900 cases are evident. From these 900 variances, we identified six strong performance trends among survey designs and populations. Confidence intervals, given with their derived methods of estimation in Appendix S2, confirm that these precision differences are highly significant for the 10,000 iterations of simulation survey run for each of the 900 cases. We summarize these survey design variance (or precision) comparison outcomes in Results subsections to follow.

For three spatial point populations (mapped in A, K and U of Fig 1 and Figs S1.1, S1.7, S1.13 in Appendix S1), complete spatial randomness [28] was achieved in practice by specifying a very short autocorrelation *range* of 1 mm. For these random (unclustered) populations, all survey designs yielded very low and effectively equal sampling error variances (*range* = 0.001 m in Figs S3.1-S3.3). Much larger variances in the estimate of density and greater differences among designs were observed in populations with wider spatial autocorrelation (*range* ≥ 5 m in Figs S3.1-S3.3), characterized by organisms clustering in broader-scale patches.

In the first subsection to follow, we evaluate the precision effect of elongating sample plot shape and specifically compare long narrow 1:100 transects with square 10:10 quadrats. Second, we compare systematic with simple random plot allocation, focusing on the effects of increasing sample plot number and varying the autocorrelation scale (and so also spatial extent) of population clustering. Third, we compare the best and worst performing designs. Summary statistics of these design performance comparisons are given in Table 1.

**Table 1.**
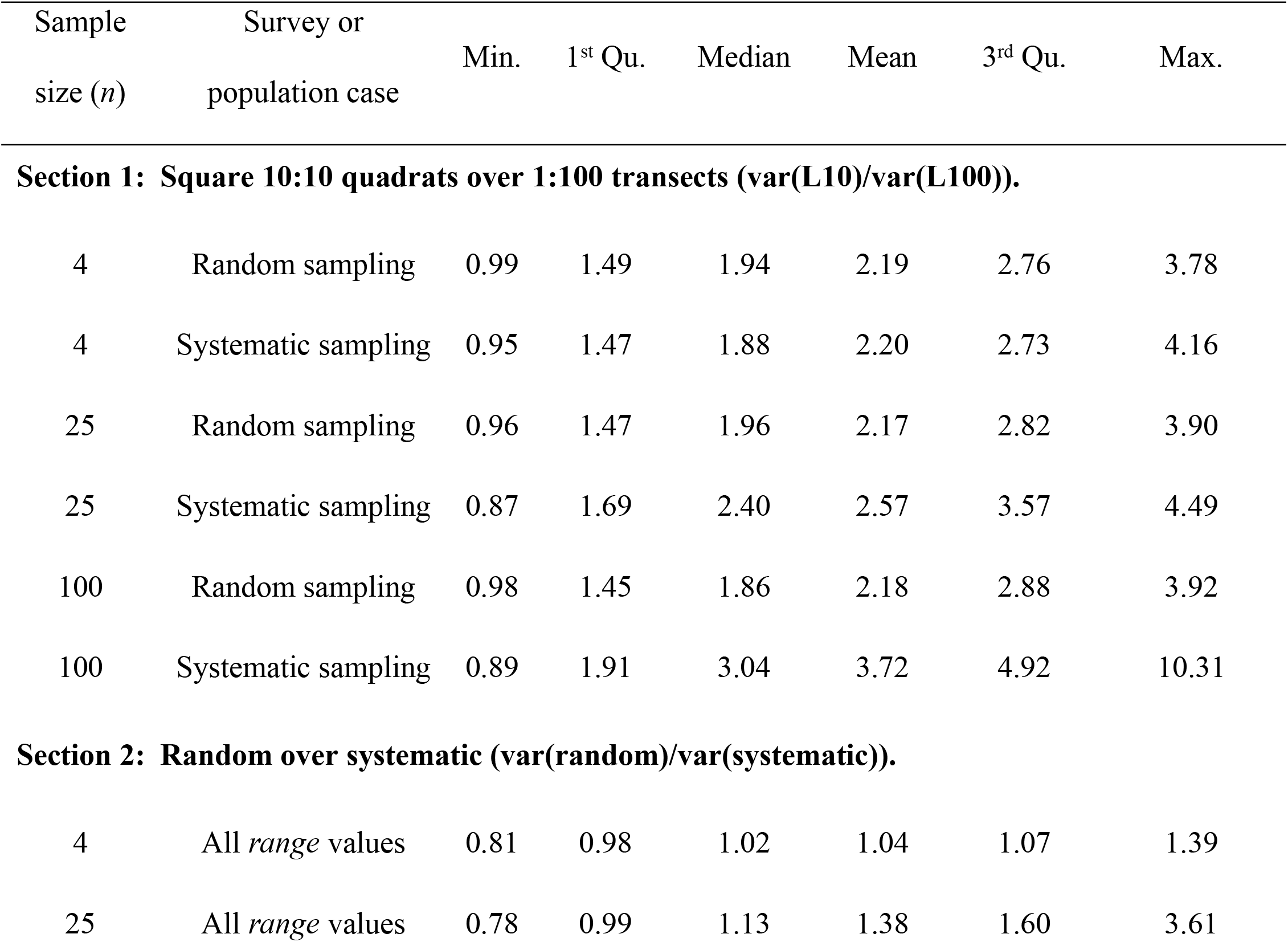
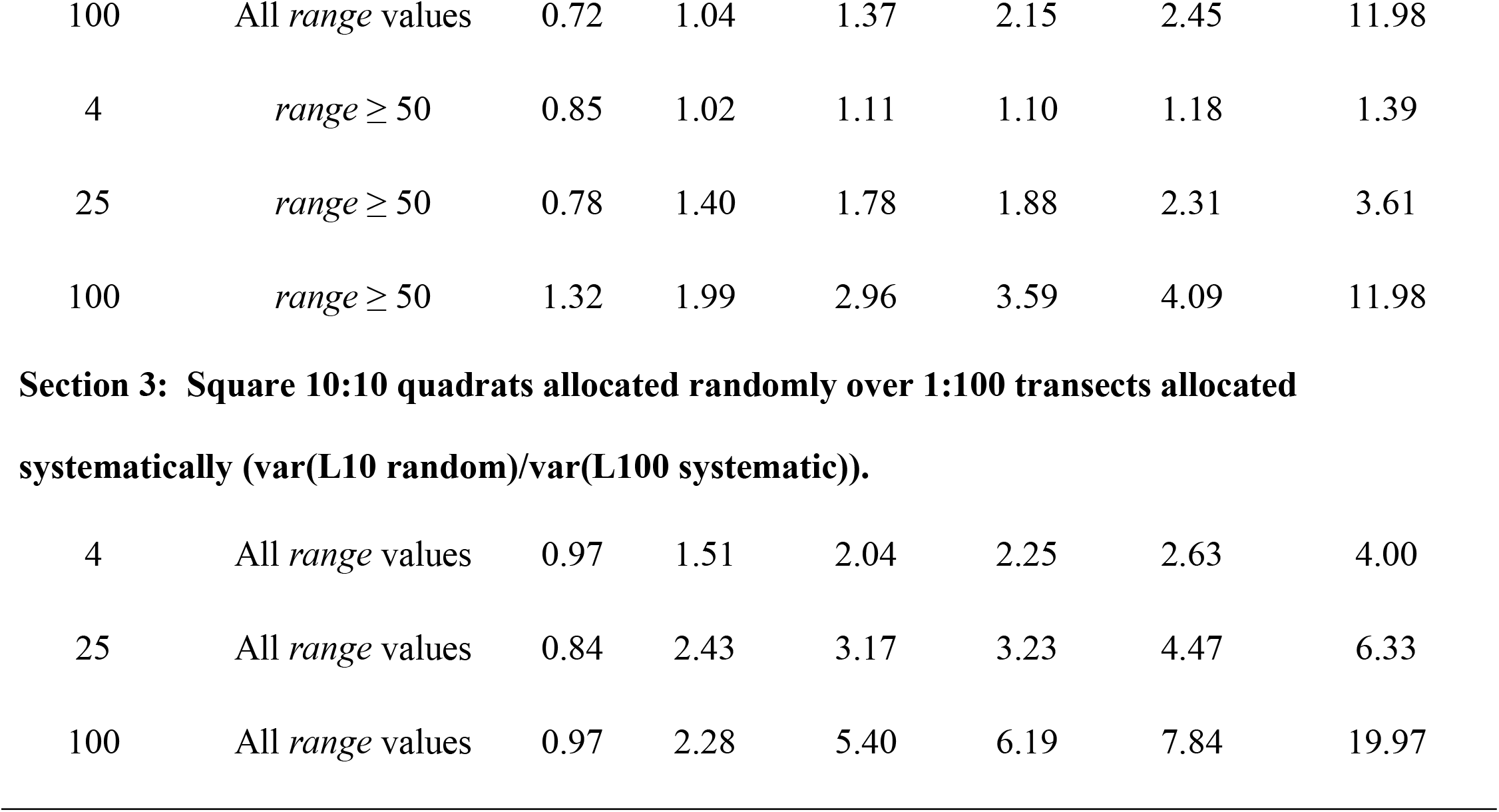
Variance ratio statistics for selected groupings of survey design and population. Section 1: square 10:10 quadrats (L10) over long narrow 1:100 transects (L100) (summarizing variance ratios shown in Fig 3); Section 2: random over systematic sample allocation (variance ratios of Fig 4); Section 3: least precise design (random square 10:10 quadrats) over most precise (systematic 1:100 transects) (variance ratios of Fig 6). Statistics of variance ratios were taken across all cases specified by the first two columns each row. In Sections 1 and 3, statistics are taken across all populations, that is, all *psill* and *range* values; in Section 2, across all *psill* values and plot shapes, for the two *range* categories shown.

### Elongated versus square sample plot shape

Elongated sample plot shapes were more precise for all clustered populations, and for most, substantially so. The most elongated (1:100) transect plot shape (L100) yielded the highest survey precision as the smallest sampling error variances (L100 blue bars smallest among every group of five plot shapes in Figs S3.1-S3.3, except for spatially random populations of *range* = 0.001). The square (10:10) plot shape (L10) nearly always produced the widest sampling error variances (red bars in Figs S3.1-S3.3) and so was least precise. Within each group of five plot shapes, elongating the plot shape (right to left, or red to blue in Figs S3.1-S3.3) nearly always produced higher survey precision shown as decreasing sampling error variances. Elongated plot shapes yielded the smallest sampling error variances for both random and systematic plot allocation, for all three sample sizes, and for all clustered populations (*range* ≥ 1-2 m).

We quantify the precision comparison of square 10:10 quadrats with 1:100 transects by the ratios of their sampling error variances (var(L10)/var(L100)). This variance ratio was computed for each combination of population, sample size, and random or systematic allocation (Fig 3). Three outcomes were evident: (i) L100 transects gave higher precision than L10 square quadrats for every non-random population examined (all bars exceeding 1 in Fig 3 except for *range* = 0.001)). Averaging over all 30 populations, for six groupings of random or systematic allocation and sample size (Table 1, Section 1), surveys using elongated transects were means of 2.2 to 3.7 times more precise than those using square quadrats. (ii) An elongated plot often yielded greater precision gains with systematic than random sampling (blue bars more often higher than red bars in Fig 3). (iii) The precision advantage of elongated (1:100) transects was greater in populations with middle-*range* autocorrelation lengths of 2 to 50 m, overlapping with the dimensions of the survey plot shapes tested. Overall, transects with dimensions of 1:100 were typically 2-3 times more precise than square 10:10 quadrats (Fig 3; Table 1, Section 1).

**Fig 3.**
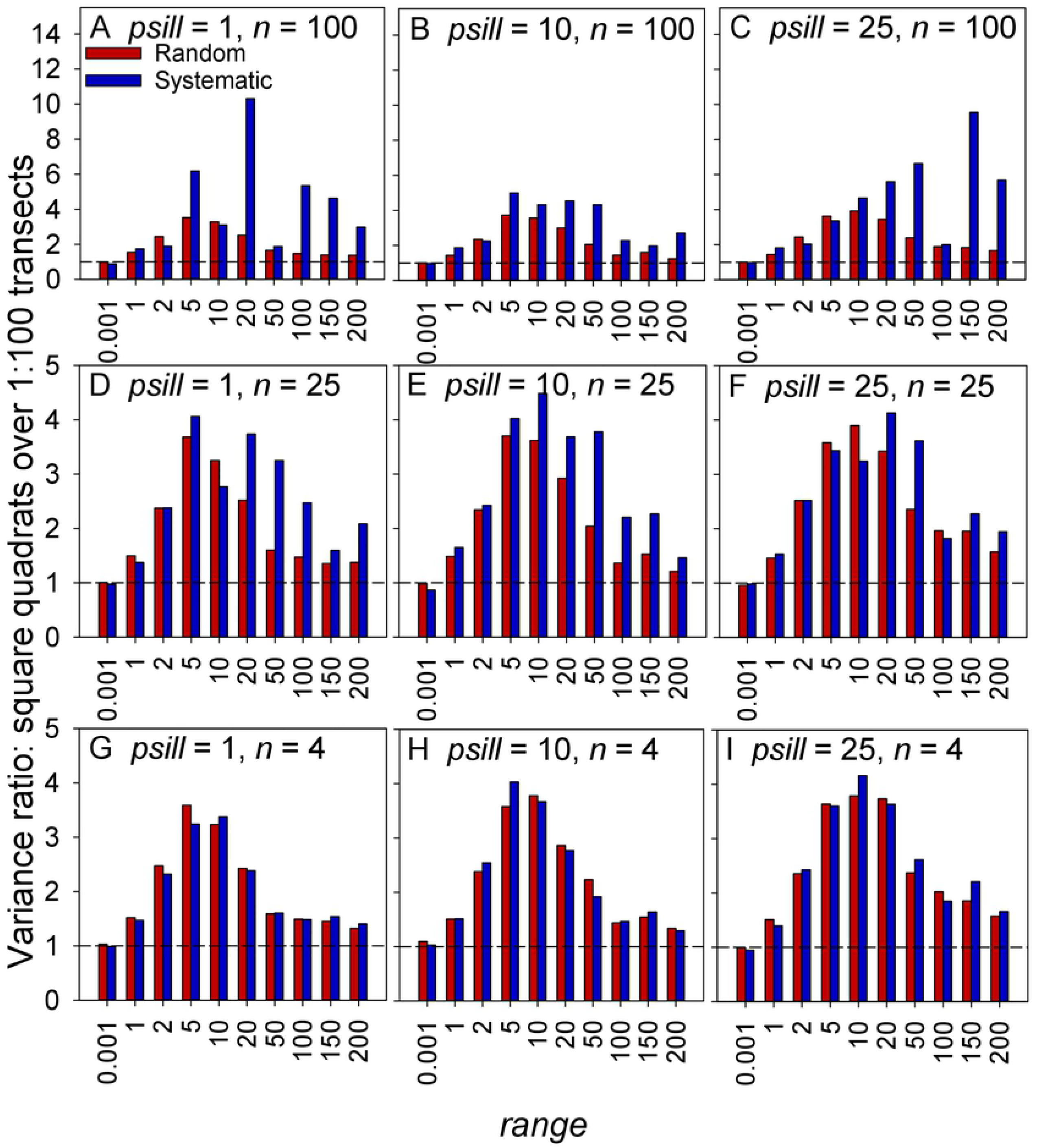
Variance ratios for assessing the relative precision of two sample plot shapes: square 10:10 quadrats (denoted L10) versus elongated 1:100 transects (L100). Sampling error variance ratios (var(L10)/var(L100)) are shown as bar heights for each combination of random (red bars) or systematic (blue bars) allocation, of survey sample size *n* (row of graphs), and of population clustering parameters *range* (x-axis) and *psill* (column of graphs). Each plotted variance ratio was computed as the sample variance of 10,000 survey mean density estimates using 10:10 quadrats divided by the sample variance of 10,000 survey mean densities using 1:100 transects. Values > 1 (bars above the dashed horizontal line) imply higher precision by the elongated plot shape.

### Systematic versus random sampling

To compare survey precision for two methods of allocating sample plots, (simple) random and (one-start aligned) systematic, we graphed the ratios of their sampling error variances as (var(random)/var(systematic)), one variance ratio for each combination of population parameters *psill* and *range*, and of survey design sample size and plot shape (Fig 4). Most random-over-systematic variance ratios being either equal to or exceeding 1 (bars near to or greater than 1 in Fig 4) imply equal or higher precision by the systematic design. Where systematic precision was higher, that was generally evident for all tested plot shapes (all bar colors in Fig 4), but the precision gain under a systematic allocation was highest for elongated transects (L100 and L50).

**Fig 4.**
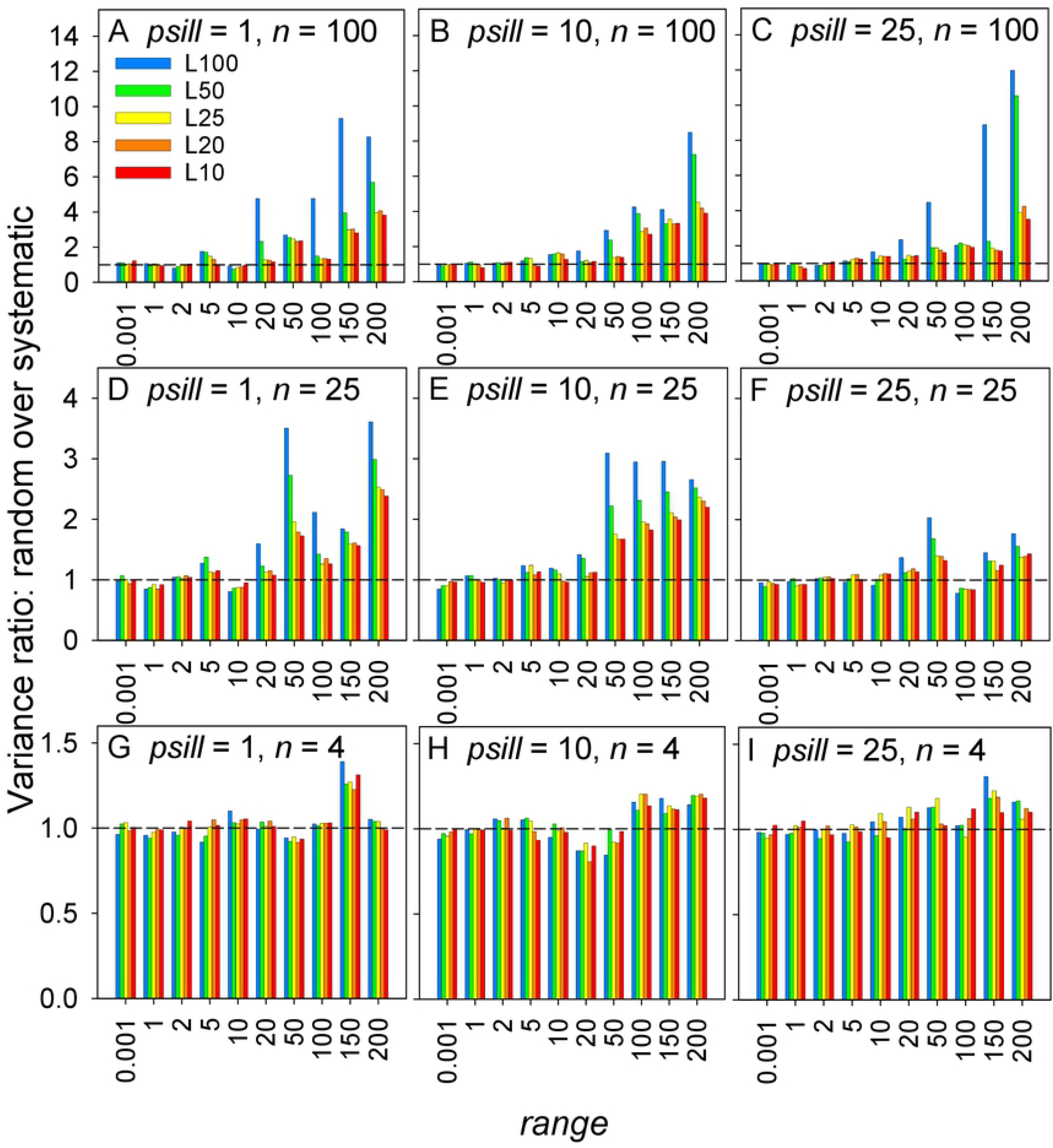
Variance ratios for assessing the relative precision of random versus systematic sampling. A variance ratio is shown for each combination of sample size *n* (row of graphs) and sample plot shape (bar color), in each of the 30 populations specified by *range* (x-axis) and *psill* (column of graphs). Each sampling error variance ratio (var(random)/var(systematic)) was computed as the sample variance of 10,000 mean densities from surveys using simple random sampling divided by the variance of 10,000 survey mean densities using (one-start aligned) systematic sampling. Values exceeding 1 imply higher precision by systematic than simple random plot allocation.

Previous studies have identified systematic designs as being more precise. However, we observed two previously unreported results that refine this general trend: (i) higher systematic precision was much more strongly evident for the larger sample sizes among those evaluated here, and (ii) systematic allocation yielded better precision than random sampling only in populations distributed with longer-range spatial autocorrelation (broader clusters). We examine these two outcomes in the next two subsections.

#### Systematic precision accelerating with sample size

Random-over-systematic variance ratios in Fig 4 increased strongly with sample size, from *n* = 4 (Fig 4G-I) to *n* = 25 (Fig 4D-F) and again to *n* = 100 (Fig 4A-C). This was primarily evident for elongated plot shapes (L100 and L50) and longer autocorrelation *range* ≥ 20-50. Means, medians and quartiles of the random-over-systematic variance ratios (Table 1, Section 2) all increased with sample size, for all *range* values and more strongly for *range* ≥ 50. Thus, systematic precision increased more rapidly with *n* than random precision. Knowing that random precision increases linearly with *n* as (*s* ^2^/*n*)^−1^, this result indicates that systematic precision increases faster than linearly with sample size.

To examine this result directly, we plotted survey precision against sample size as number of sample plots (*n*). We separately consider both random and systematic allocations (Fig 5). 18 test populations were examined, six levels of autocorrelation *range* (0.001, 5, 20, 50, 100, 200 m) and all three levels of *psill* (1, 10, 25). As explained in Methods, selecting two plot shapes, L50 (2:50 transects, Fig 5A) and L10 (square quadrats, Fig 5B), permitted the inclusion for systematic designs of a fourth sample size, *n* = 16.

**Fig 5.**
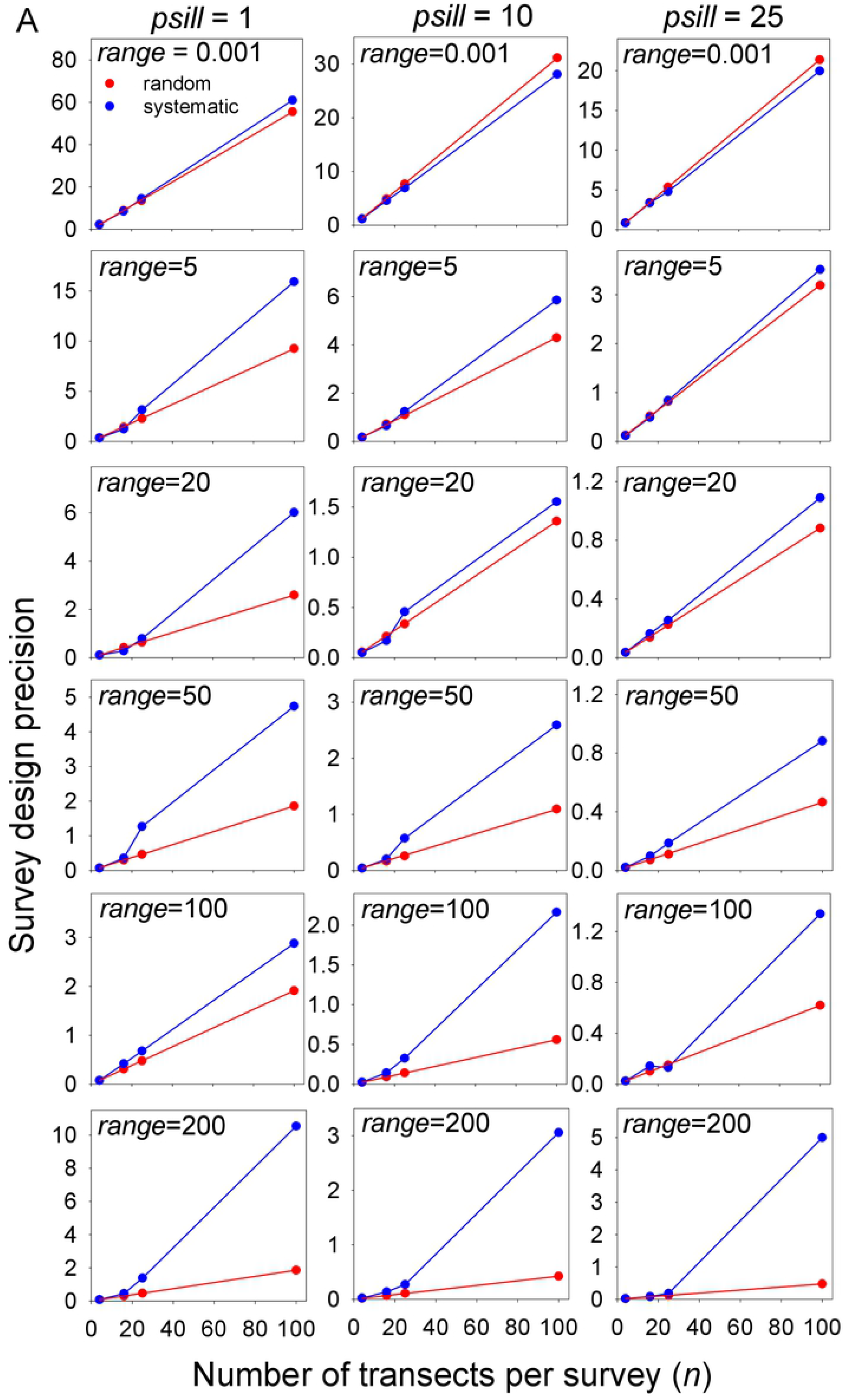
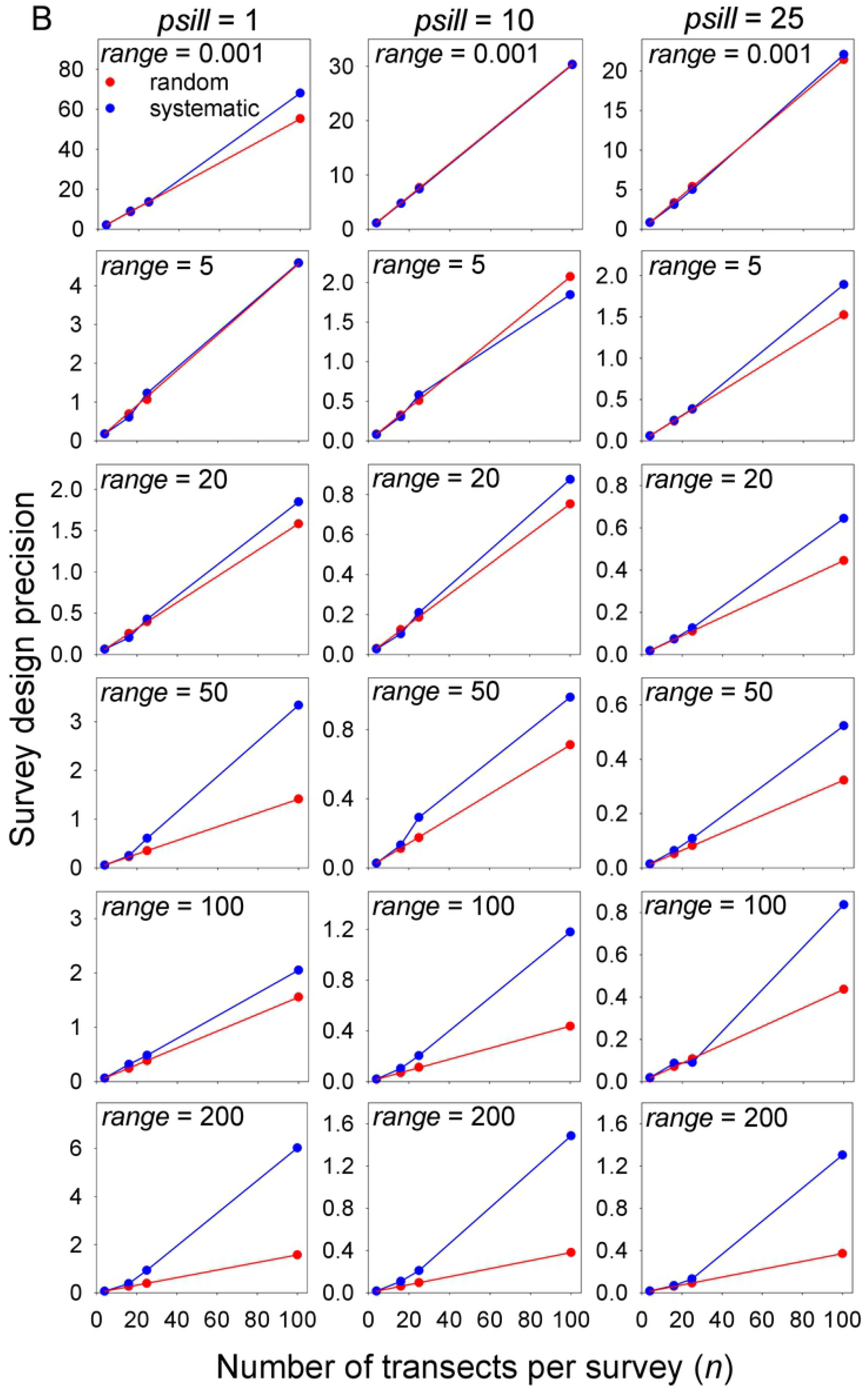
Survey precision versus sample size for random and systematic sample allocations. Precision, computed as the inverse of sampling error variance, is plotted against sample size (*n* = 4, 16, 25 and 100 sample plots) in each of 18 simulated populations. Two plot shapes were used: (panel A) a 2:50 transect plot shape (L50) and (panel B) a 10:10 square quadrat (L10). The red lines passing through the random design precision points are ordinary least squares regressions. The blue lines plotted for the systematic design directly connect the computed precision estimates shown.

The trend for systematic precision dependence on sample size (blue lines in Fig 5) confirms non-linear, accelerating dependence, here notably for *n* ≥ 25. The rate of this acceleration increased strongly with longer autocorrelation *range* (greater divergence of blue plots from red plots in descending rows of graphs in Fig 5). As basic statistics predicts, the precision of random sampling increased strictly linearly with sample size (red lines in all graphs of Fig 5). Overall, a more rapid increase of systematic than random precision, systematic precision accelerating with increasing number of sample plots, was evident for both 2:50 transects (Fig 5A) and square quadrats (Fig 5B). We could find no previous report of systematic precision increasing faster than linearly with sample size.

#### Dependence on population spatial autocorrelation length

Sampling error variances depended strongly on the length scale of population clustering, controlled here by the *range* parameter of spatial autocorrelation. Increasing the *range* produces populations with correspondingly wider clusters and broader areas of zero density (see Fig 1; finer detailed mapping of spatial clusters are shown in Figs S1.1-S1.6, S1.7-S1.12, S1.13-S1.18). For all survey designs and *psill* values (all graphs in Figs S3.1-S3.3), sampling error variances increased steeply with population *range* up to about 20 or 50 m. Trends varied for *range* > 50.

For *range* ≤ 10, random-over-systematic variance ratios are all near 1 (bar heights ∼ 1 in Fig 4), implying similar precision by systematic and random sampling. Higher precision by the systematic design (Fig 4 bar heights > 1) was evident primarily for *range* ≥ 20-50. For *n* = 100 and *range* ≥ 50, averaging over all plot shapes and *psill* values, a random-over-systematic variance ratio mean of 3.59 (Table 1, Section 2) implies a systematic-versus-random precision advantage of 3-to 4-fold. For samples of *n* = 25, with *range* ≥ 50 (Fig 4D-F), mean sampling error variances were 1.88 times wider for random than systematic sampling (Table 1, Section 2). Thus, in addition to the strong effect of sample size described above, systematic designs yielded substantially higher precision than random designs only in populations with longer-range spatial autocorrelation (*range* ≥ 20 or 50 in Fig 4), i.e. only in populations with broader clusters.

In contrast, systematic-versus-random precision showed no clear trend under variation of the other population clustering parameter, *psill*, which controls the maximum density within clusters. Comparing populations of increasing *psill* (graphs from left to right in each row of Fig 4), random-over-systematic variance ratios did not consistently rise or fall.

In summary, allocating samples systematically delivered more precise estimates of population density in clustered populations, but only for *n* ≥ 25 sample plots and in populations with longer-range clustering (*range* ≥ 20-50). This population autocorrelation *range* of ∼ 20-50 m overlaps with the spread of tested survey sample plot lengths, from 10 to 100 m, suggesting an interaction of population cluster size and sample plot dimension. A stronger and much more consistent effect on the precision advantage of systematic over random sampling was its dependence on sample size.

Averaged across all *psill* values and all plot shapes (Table 1, Section 2), for *n* = 25, a systematic design improved survey precision by mean factors of 1.38 (all *range* values) and 1.88 (*range* ≥ 50) and, for *n* = 100, by factors of 2.15 (all *range* values) and 3.59 (*range* ≥ 50).

### Systematic long narrow transects versus random quadrats

Finding that elongated plot shapes yielded higher precision, as did systematic allocation for many cases, we compared the most precise survey design (systematically allocated 1:100 transects) with the least precise (randomly allocated square quadrats). Computing their variance ratios (var(L10 random)/var(L100 systematic)), systematic elongated transects yielded higher precision than random quadrats for every combination of sample size, *psill*, and *range* (Fig 6, all ratios > 1 except for *range* = 0.001). Elongated transects allocated systematically were approximately 2 times (for *range* ∼ 2 in Fig 6) ranging up to 20 times (for *n* = 100, *psill* = 25, *range* = 200 in Fig 6C) more precise than random square quadrats. Averaging across all populations yielded mean values for these sampling error variance ratios of 2.25 for *n* = 4, 3.23 for *n* = 25, and 6.19 for *n* = 100 (Table 1, Section 3). These averages confirm large (doubling) to very large (five-fold or more) gains in efficiency for designs employing elongated systematic transects.

**Fig 6.**
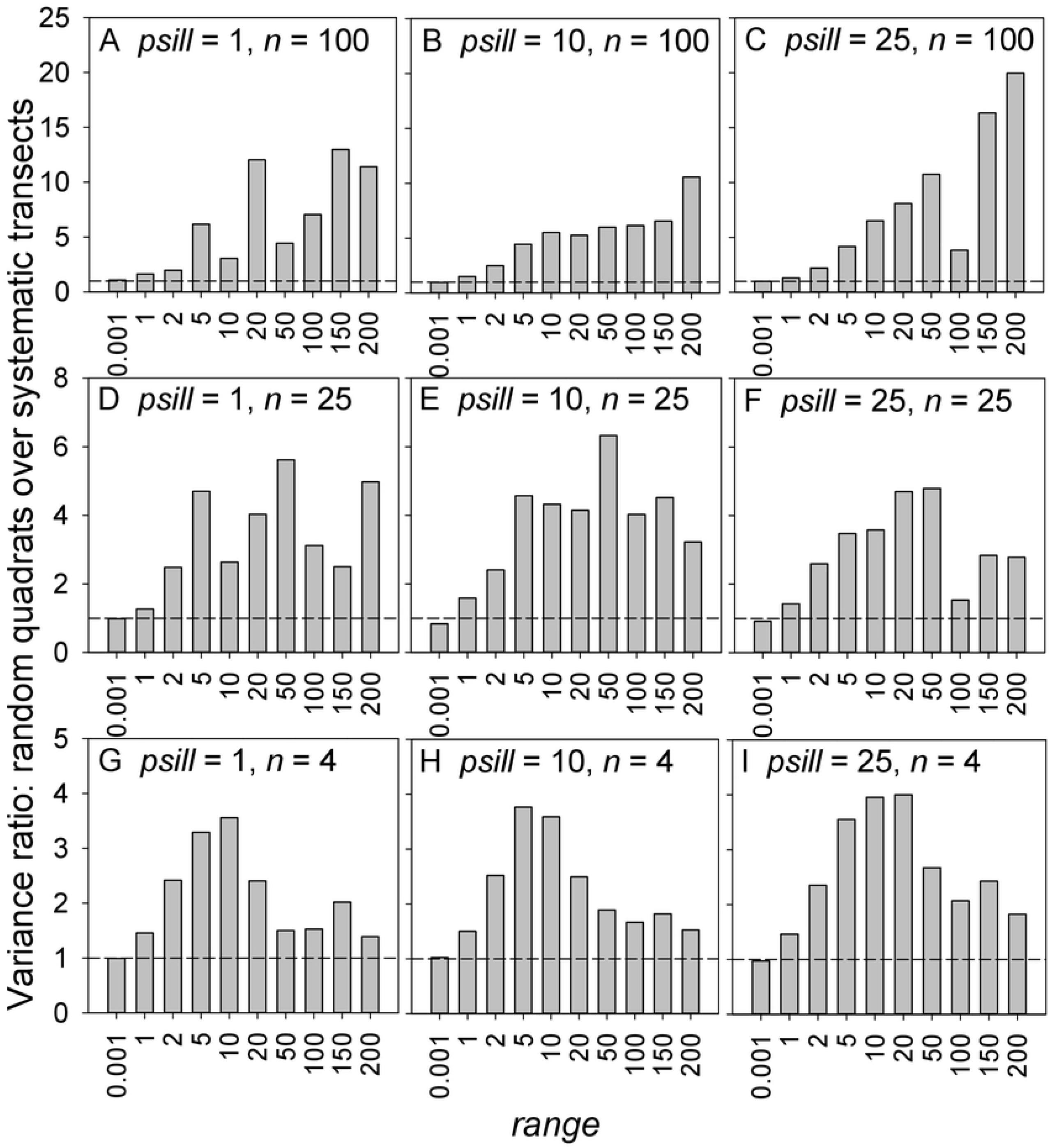
Variance ratios for assessing the relative precision of random square quadrats versus systematic 1:100 transects. A variance ratio was computed for each combination of *psill* (column of graphs), *range* (x-axis) and survey sample size *n* (row of graphs) as the sample variance of 10,000 survey mean densities using randomly allocated 10:10 quadrats, divided by the sample variance of 10,000 survey mean densities using systematically allocated 1:100 transects. Values > 1 imply higher precision for systematic elongated transects.

## Discussion

Our aim was to evaluate the performance of sample plot survey designs, focusing on plot shape, in a wide range of clustered populations. We report the following performance outcomes: (i) Elongating the sample plot shape produced higher precision in all clustered populations, for both systematic and random allocations, and for all three sample sizes we evaluated. A long thin shape of dimensions 1:100 produced precisions averaging 2-3 times higher than a square shape across the 30 populations tested. (ii) The precision of systematic plot allocation increased faster with sample size than the linear increase of random precision, accelerating with sample plot number. This new and unanticipated (here purely computational) outcome strongly favors systematic designs as sample sizes increase with automated measuring technologies and remote sensing. This result, of rapid increase in systemic precision with sample plot number, warrants further, possibly analytic, statistical investigation. (ii.a) This survey precision acceleration showed a strong increasing dependence on population autocorrelation range. As previously known, systematic plot allocation showed overall higher precision than simple random allocation, but we found this improvement (iii) only for surveys with ≥ 25 sample plots, and (iv) only in populations with longer-range spatial autocorrelation, notably where the breadth of spatial clusters are similar to or greater than sample plot dimensions. (v) However, systematic-versus-random precision showed no consistent dependence on the maximum density within clusters. Outcomes iii-v suggest rules of thumb identifying where a systematic design should be expected to improve the precision of population sampling. (vi) An elongated plot shape combined with a systematic allocation showed the highest precision, giving mean improvements relative to random square sample plots of approximately 200% (*n* = 4), 300% (*n* = 25), and 600% (*n* = 100).

These results confirm that counting organisms within more elongated plot shapes that stretch across broader reaches of habitat and (sometimes) spreading sample plots evenly across the study region capture a more representative sample of mean density in clustered populations. An additional design question is whether more but smaller plots, still maintaining constant area searched, would further improve survey precision. Previous studies [4, 19, 20, 37] have concluded it would.

In addition to more precise estimates of mean population density, long narrow transects can be subdivided to provide geostatistical data for mapping the spatial distribution of population density within a study region. Recording organism sub-counts in, say 1- or 2-m quadrats sequentially along 100-m transects, provides detailed information on the spatial autocorrelation in population density over shorter spatial scales [38]. Measured autocorrelation over shorter distance underpins both kriging (as the estimated nugget and the semivariogram at shorter-range separation distances) and likelihood-based geostatistical inference (in which the autocorrelation function is employed directly [39, 40]_)_. A line sequence of counts is also the input data used in spatial clustering pattern identification [41], including estimation of cluster radius by local quadrat variance methods [42-45]. An alternative proposed for measuring shorter-distance autocorrelation is to add a second set of supplementary sample plots at short distances away from the set of primary samples [40, 46-48]. However, when they are feasible, subplots within (along) each transect avoids the need for supplementary samples with attendant survey cost savings since no additional habitat needs to be covered, will produce many more short- and medium-distance sample pairs for computing autocorrelation versus distance, and is simpler to implement by avoiding the mathematical and statistical elaboration required to avoid bias in supplemented-sample designs.

An elongated plot shape, to provide a more representative sample of the overall mean, can be aligned parallel to known habitat gradients, further improving density estimate precision. If autocorrelation varies in two dimensions and measuring that anisotropic autocorrelation is an objective [49], directional variation in spatial autocorrelation can also be quantified using subplots along elongated transects of systematically or randomly varied directional orientation which should not bias the estimates of overall mean population density. In field practice, researchers moving along a transect can in some situations more easily identify the habitat to follow, cover and search. A line can be deployed (e.g. by rope or laser) and organisms counted that fall within a fixed distance from the line [6].

Systematic sample allocation has long been found to be more precise in the presence of autocorrelation for both one-dimensional [35, 50] and two-dimensional populations [36, 51-54], including many in an ecological context [34, 55-58]. One important drawback of a systematic design is that, in practice from a single systematic survey sample, no formula or algorithm has yet been derived to accurately estimate its sampling error variance [59-61], which is often over-estimated by the standard formula *s*^2^/*n* [34, 54, 62] that accurately estimates the variance of the mean under random sampling. Since systematic sampling nearly always has smaller or equal variance than random sampling, confidence intervals obtained using *s*^2^/*n* are conservative.

However, this inability to reliably estimate systematic variance does not affect the choice of plot shape; *s*^2^/*n* applies for random allocation of any plot shape. In these simulations, the 1/*n* dependence of random sample variance is shown by the exact linearity of precision versus sample size evident for both plot shapes used, that is in both panels of Fig 5, square quadrats (red lines of Fig 5B) and 2:50 transects (red lines of Fig 5A).

In multi-species surveys that use sample plots, for conservation monitoring or sampling whole food webs, these (and other) survey design rules of thumb can be usefully applied to all species counted. In choosing sample plot dimensions, consideration should be given to the spatial autocorrelation properties of targeted species [19].

The design comparisons above show that sampling error variance ratios provide a direct and easily applied statistic for power analysis when comparing two specific designs, one of which is random. With random sampling error variance accurately estimated by *s*^2^/*n*, increasing sample plot number *n* by any factor reduces random survey sample variance by the same factor.

The average variance ratio of 6 (600%) we observed for 100 random square quadrats compared with 100 systematic elongated transects implies that a field researcher would need, on average, to obtain counts from 6 times more (*n* = 600) randomly allocated square (10:10) quadrats, and thus would need to cover 6 times more habitat, to achieve the precision of *n* = 100 equal-area elongated (1:100) systematic transects.

More generally, long thin transects yielded superior precision for both systematic and random allocations. When a simple random allocation is preferred, a 1:100 versus a 10:10 plot shape more than doubled survey precision, and so also its efficiency, averaged among the 30 clustered populations we tested (Table 1).

## Conclusions

Resources for conservation monitoring are limited and the need is growing to monitor natural populations in the face of climate change and other anthropogenic impacts on earth ecosystems. The 600% average efficiency advantage we found for 100 systematic transects compared with 100 random quadrats is unexpectedly large. A choice of a long narrow plot shape, regardless of allocation method or sample size, more than doubled survey efficiency. If opportunities for these efficiency gains could be realized in practice, it would produce more statistically informative field measurements of population change and of absolute population density on given research budgets. These would more confidently identify statistical correlates of change in threatened populations at risk, permitting better targeted and more effective management decision making.

## Acknowledgments

We thank Jason Tanner, Craig Noell, Zoe Doubleday, Peter Diggle, and Marie-Josée Fortin who provided valuable comments on the draft manuscript.

## Supporting information

**Appendix S1**: Pixelated maps showing density in 1-m^2^ tiles for 18 of the 30 generated test populations (Fig 1).

**Appendix S2**: Statistical formulas applied for estimating confidence intervals of survey sampling error variance, precision, and variance ratio.

**Appendix S3**: Bar plots of computed sampling error variances from simulated surveys for all 900 combinations of sample plot shape, random or systematic plot allocation, sample size, and population autocorrelation parameters *range* and *psill*.

**Dataset S1**: R script file used to generate spatially autocorrelated populations

**Dataset S2**: R script file containing functions used to simulate systematic and random sampling of these populations for differing plot shapes and sample sizes.

## References

1. Field SA, Tyre AJ, Possingham HP. Optimizing allocation of monitoring effort under economic and observational constraints. J Wildl Manage. 2005;69(2): 473–482.

2. Legendre P, Fortin M-J. Spatial pattern and ecological analysis. Vegetatio. 1989;80(2): 107–138.

3. Dungan JL, Perry JN, Dale MRT, Legendre P, Citron-Pousty S, Fortin M-J, et al. A balanced view of scale in spatial statistical analysis. Ecography. 2002;25(5): 626–640.

4. Pennington M, Volstad JH. Assessing the effect of intra-haul correlation and variable density on estimates of population characteristics from marine surveys. Biometrics. 1994: 725–732.

5. Pennington M. Estimating the mean and variance from highly skewed marine data. Fish Bull. 1996;94: 498–505.

6. McGarvey R, Mayfield S, Byth K, Saunders T, Chick R, Foureur B, et al. A diver survey design to estimate absolute density, biomass, and spatial distribution of abalone. Can J Fish Aquat Sci. 2008;65(9): 1931–1944.

7. Kotwicki S, De Robertis A, Ianelli JN, Punt AE, Horne JK. Combining bottom trawl and acoustic data to model acoustic dead zone correction and bottom trawl efficiency parameters for semipelagic species. Can J Fish Aquat Sci. 2013;70(2): 208–219.

8. Hamner WM, Carleton JH. Copepod swarms: attributes and role in coral reef ecosystems. Limnol Oceanogr. 1979;24(1): 1–14.

9. Pepin P, Helbig JA. Sampling variability of ichthyoplankton surveys—exploring the roles of scale and resolution on uncertainty. Fish Res. 2012;117: 137–145.

10. Vandermeer J, Perfecto I, Philpott SM. Clusters of ant colonies and robust criticality in a tropical agroecosystem. Nature. 2008;451(7177): 457–459.

11. Greig-Smith P. Quantitative plant ecology. 3rd ed: University of California Press; 1983.

12. Hayek LAC, Buzas MA. Surveying natural populations: quantitative tools for assessing biodiversity: Columbia University Press; 2010.

13. Martin TG, Wintle BA, Rhodes JR, Kuhnert PM, Field SA, Low-Choy SJ, et al. Zero tolerance ecology: improving ecological inference by modelling the source of zero observations. Ecol Lett. 2005;8(11): 1235–1246.

14. Christidis BG. The importance of the shape of plots in field experimentation. J Agric Sci. 1931;21(1): 14–37.

15. Lyon TL. Some experiments to estimate errors in field plat tests. Proc Am Soc Agron. 1911;3(1): 89–114.

16. Mercer WB, Hall AD. The experimental error of field trials. J Agric Sci. 1911;4(2): 107–132.

17. Clapham AR. The form of the observational unit in quantitative ecology. J Ecol. 1932: 192–197.

18. Bormann FH. The statistical efficiency of sample plot size and shape in forest ecology. Ecology. 1953;34(3): 474–487.

19. Kenkel NC, Juhász-Nagy P, Podani J. On sampling procedures in population and community ecology. Vegetatio. 1989;83(1): 195–207.

20. Justesen SH. Influence of size and shape of plots on the precision of field experiments with potatoes. J Agric Sci. 1932;22(2): 366–372.

21. Kalamkar RJ. Experimental error and the field-plot technique with potatoes. J Agric Sci. 1932;22(2): 373–385.

22. Pechanec JF, Stewart G. Sagebrush-grass range sampling studies: size and structure of sampling unit. J Am Soc Agron. 1940;32: 669–82.

23. Wight JR. The sampling unit and its effect on saltbush yield estimates. J Range Manage. 1967;20(5): 323–325.

24. Van Dyne GM, Vogel WG, Fisser HG. Influence of small plot size and shape on range herbage production estimates. Ecology. 1963;44(4): 746–759.

25. Papanastasis VP. Optimum size and shape of quadrat for sampling herbage weight in grasslands of northern Greece. J Range Manage. 1977;30(6): 446–449.

26. Brummer JE, Nichols JT, Engel RK, Eskridge KM. Efficiency of different quadrat sizes and shapes for sampling standing crop. J Range Manage. 1994;47(1): 84–89.

27. Cochran WG. Sampling Techniques. 3rd ed. New York: John Wiley & Sons; 1977.

28. Diggle PJ. Statistical analysis of spatial and spatio-temporal point patterns. 3rd ed: Chapman and Hall; 2013.

29. Wagner HH, Holderegger R, Werth S, Gugerli F, Hoebee SE, Scheidegger C. Variogram analysis of the spatial genetic structure of continuous populations using multilocus microsatellite data. Genetics. 2005;169(3): 1739–1752.

30. Ciannelli L, Fauchald P, Chan K-S, Agostini VN, Dingsør GE. Spatial fisheries ecology: recent progress and future prospects. J Marine Syst. 2008;71(3-4): 223–236.

31. Goslee SC. Behavior of vegetation sampling methods in the presence of spatial autocorrelation. Plant Ecol. 2006;187(2): 203–212.

32. Pebesma EJ. Multivariable geostatistics in S: the gstat package. Comput Geosci. 2004;30(7): 683–691.

33. R Core Team. R: A language and environment for statistical computing: R Foundation for Statistical Computing, Vienna, Austria; 2015. Available from: http://www.R-project.org/.

34. McGarvey R, Burch P, Matthews JM. Precision of systematic and random sampling in clustered populations: habitat patches and aggregating organisms. Ecol Appl. 2016;26(1): 233–248.

35. Madow WG, Madow LH. On the theory of systematic sampling, I. Ann Math Stat. 1944;15(1): 1–24.

36. Quenouille MH. Problems in plane sampling. Ann Math Stat. 1949;20: 355–375.

37. Pennington M, Volstad JH. Optimum size of sampling unit for estimating the density of marine populations. Biometrics. 1991;47: 717–723.

38. McGarvey R, Feenstra JE, Mayfield S, Sautter EV. A diver survey method to quantify the clustering of sedentary invertebrates by the scale of spatial autocorrelation. Mar Freshwater Res. 2010;61(2): 153–162.

39. Diggle PJ, Ribiero PJ. Model-based geostatistics: Springer-Verlag; 2007.

40. Diggle P, Lophaven S. Bayesian geostatistical design. Scand J Stat. 2006;33(1): 53–64.

41. Dale MRT, Dixon P, Fortin MJ, Legendre P, Myers DE, Rosenberg MS. Conceptual and mathematical relationships among methods for spatial analysis. Ecography. 2002;25(5): 558–577.

42. Hill MO. The intensity of spatial pattern in plant communities. J Ecol. 1973;61: 225–235.

43. Dale MRT. Spatial pattern analysis in plant ecology Cambridge University Press; 1999.

44. Perry JN, Liebhold AM, Rosenberg MS, Dungan J, Miriti M, Jakomulska A, et al. Illustrations and guidelines for selecting statistical methods for quantifying spatial pattern in ecological data. Ecography. 2002;25(5): 578–600.

45. Leeworthy G. Application of the two-term local quadrat variance analysis in the assessment of marine invertebrate populations: preliminary findings on the sea cucumber Actinopyga echinites. SPC Beche de Mer Information Bulletin. 2007;26: 26–30.

46. Lark RM. Optimized spatial sampling of soil for estimation of the variogram by maximum likelihood. Geoderma. 2002;105(1-2): 49–80.

47. Müller WG, Zimmerman DL. Optimal designs for variogram estimation. Environmetrics. 1999;10(1): 23–37.

48. Chipeta M, Terlouw D, Phiri K, Diggle P. Inhibitory geostatistical designs for spatial prediction taking account of uncertain covariance structure. Environmetrics. 2017;28(1): e2425.

49. Fortin M-J. Effects of sampling unit resolution on the estimation of spatial autocorrelation. Ecoscience. 1999;6(4): 636–641.

50. Cochran WG. Relative accuracy of systematic and stratified random samples for a certain class of populations. Ann Math Stat. 1946;17(2): 164–177.

51. Bellhouse DR. Some optimal designs for sampling in two dimensions. Biometrika. 1977;64(3): 605–611.

52. Yates F. Systematic sampling. Philos Trans R Soc Lond A. 1948;241(834): 345–377.

53. Das AC. Two dimensional systematic sampling and the associated stratified and random sampling. Sankhyā. 1950: 95–108.

54. Matérn B. Spatial Variation, volume 36 of Lecture Notes in Statistics. Berlin: Springer-Verlag; 1986.

55. Dunn R, Harrison AR. Two-dimensional systematic sampling of land use. J R Stat Soc Ser C Appl Stat. 1993;42(4): 585–601.

56. Ambrosio L, Iglesias L, Marin C, Del Monte J. Evaluation of sampling methods and assessment of the sample size to estimate the weed seedbank in soil, taking into account spatial variability. Weed Res. 2004;44(3): 224–236.

57. Fewster RM. Variance estimation for systematic designs in spatial surveys. Biometrics. 2011;67(4): 1518–1531.

58. Aune-Lundberg L, Strand G-H. Comparison of variance estimation methods for use with two-dimensional systematic sampling of land use/land cover data. Environ Modell Softw. 2014;61: 87–97.

59. Bellhouse DR, Sutradhar BC. Variance estimation for systematic sampling when autocorrelation is present. The Statistician. 1988;37(3): 327–332.

60. D’Orazio M. Estimating the variance of the sample mean in two-dimensional systematic sampling. J Agric Biol Environ Stat. 2003;8(3): 280–295.

61. Wolter KM. Introduction to variance estimation. 2nd ed. New York: Springer Science + Business Media; 2007.

62. McBratney AB, Webster R. How many observations are needed for regional estimation of soil properties? Soil Sci. 1983;135(3): 177–183.

